# Enzymatic Ligation Strategy to Enhance Electrospray Ionization Efficiency and Liquid Chromatography-Mass Spectrometry of DNA and RNA Oligonucleotides

**DOI:** 10.64898/2026.04.09.717506

**Authors:** Max D. Sharin, Nina J. Fitzgerald, Stephen M. Kennedy, Isabella G. Park, Kevin D. Clark

## Abstract

Mass spectrometry (MS) is a powerful technique for characterizing modified RNA as it directly sequences and quantifies all mass-altering modifications simultaneously. However, the physicochemical properties of RNA result in poor ionization efficiencies during electrospray ionization, presenting a major barrier to sensitive MS measurements necessary for low abundance RNA samples and RNAs with low modification stoichiometries. Here, we report a ligation-based approach to increase ionization efficiencies of RNA oligonucleotides. We show that short (∼5 nt), chemically modified DNA oligonucleotides can be enzymatically ligated to RNA to serve as MS signal enhancers. Among a series of signal enhancers appended with various alkyl and alkylimidazolium functional groups, we found that decyl-functionalized derivatives improved MS sensitivity by ∼15-fold compared to unlabeled oligonucleotide. When ligated to RNA standards, the decyl-modified signal enhancer increased MS signals 2-4-fold with the additional benefit of improved retention during liquid chromatography (LC) separations without ion pairing agents. To apply the ligation-based approach to RNase T1 digests of longer RNAs, a multi-step enzymatic approach was optimized to maximize ligation efficiencies. We then ligated signal enhancers to a yeast transfer RNA (tRNA) digest and observed increased MS signals for numerous sequence-informative digestion products. Importantly, the sequences of RNA oligonucleotides ligated to signal enhancers were readily determined by tandem mass spectrometry with collision-induced dissociation. This ligation-based strategy for enhancing LC-MS/MS characterization of RNA creates opportunities to measure low abundance RNA samples and their modifications.

## Introduction

Every organism contains a diverse collection of ribonucleic acids (RNAs) that are primarily composed of four canonical nucleosides: adenosine (A), uridine (U), guanosine (G), and cytidine (C). RNAs serve as intermediaries between heritable genetic information and proteins yet have a host of other important functions including catalysis,^1,2^ splicing,^3^ and regulating gene expression.^4^ This remarkably broad range of RNA functions is increasingly linked to the more than 170 post-transcriptional modifications of RNA that have been discovered across the domains of life.^5–7^ These modifications comprise what is known as the epitranscriptome and pattern virtually all cellular RNAs including transfer RNA (tRNA), ribosomal RNA (rRNA), and messenger RNA (mRNA). Enzymatic writers of the epitranscriptome deposit modifications that are chemically diverse ranging from base methylations and isomerizations to hypermodifications like aminocarboxypropyluridine (acp^3^U) or 5-methoxycarbonylmethyl-2-thiouridine (mcm^5^s^2^U). RNA modification profiles vary throughout development and are cell type specific,^8,9^ suggesting important regulatory effects. Functions of RNA modifications depend on sequence position and include regulating mRNA transcript stability,^10^ modulating tRNA decoding efficiency,^11,12^ and stabilizing RNA structures.^13^ These functions are potentiated by specialized reader proteins and are further modulated by enzymes that “erase” select RNA modifications.^14^ A thorough and simultaneous mapping of all post-transcriptional modifications, especially low abundance modifications which are difficult to measure, is crucial to discern their functions in various biological contexts.

There are several orthogonal approaches to characterizing modified RNAs including next generation sequencing (NGS), nanopore direct RNA sequencing (DRS), and mass spectrometry (MS). NGS approaches typically involve antibody enrichment or chemical derivatization of a select modification and have identified sites of modification transcriptome-wide.^15–19^ Although these methods leverage enzymatic amplification to afford high sensitivity, they only measure a single or few modification types per experiment and provide limited information regarding modification stoichiometry. Moreover, NGS-based approaches do not directly detect RNA modifications, increasing the likelihood of false positive results.^20,21^ Confidence in DRS measurements hinges on matching ionic current patterns to modifications, which has only been well characterized for a handful of modifications, namely *N6*-methyladenosine, 5-methylcytidine, and pseudouridine.^22^ Conversely, MS methods simultaneously and directly measure any mass altering modifications of RNA biopolymers. MS-based approaches typically involve enzymatic digestion, liquid chromatography (LC) separation and tandem mass spectrometry (MS/MS) measurements that are ideally suited for quantitative assessment of modification stoichiometries.^23,24^ However, a major limitation to LC-MS/MS methods is their notoriously poor sensitivity toward RNA, which restricts MS applications to bulk RNA samples and abundant RNA modifications.

Measuring RNA modifications by LC-MS/MS often involves LC separation of nucleolytic fragments of RNAs that are then subjected to electrospray ionization (ESI) to generate multiply charged gas phase RNA ions. The poor sensitivity of LC-MS methods for RNA is largely due to the low ionization efficiency (IE) of oligonucleotides during ESI. The process by which gas phase ions are produced by ESI can be explained by multiple theories that involve analyte partitioning to the surface of the electrospray droplet.^25–28^ The hydrophilic nature of oligonucleotides originating from the polyanionic backbone and numerous hydrogen bond donors and acceptors impedes the partitioning of RNA to the droplet surface. Thus, the physicochemical properties of nucleic acids hinder their escape from electrospray droplets as gas phase ions, resulting in low IEs and poor MS sensitivity for RNA. Although these effects can be partially mitigated by mobile phase additives like ion pairing agents^29^ or fluorinated alcohols,^30^ these reagents tend to contaminate LC-MS systems and thus require dedicated instruments that may not be suitable for general-purpose labs. Alternative LC methods include ion pair-free reversed phase^31^ and HILIC methods,^32^ yet these approaches still require large amounts of RNA sample. As a result, the poor IE of RNA remains a formidable barrier to MS detection and characterization of RNA modifications.

One successful approach to improving IEs for biomolecules involves directly altering their physicochemical properties via chemical derivatization. Extremely sensitive measurements of peptides and other biomolecules have been accomplished via derivatization strategies that enhance MS detection up to 15,000 fold,^33^ yet chemical derivatization strategies to deliberately increase oligonucleotide ionization efficiencies are lacking.^34^ For the analysis of free nucleosides or nucleotides, derivatization reagents have been employed to enhance signals from 3 to ∼1000 fold.^35–37^ Some derivatization strategies for intact RNA analysis by LC-MS include 3′ terminal phosphate labeling with a silylation reagent,^38^ and the incorporation of biotin hydrophobic end labels.^39,40^ These strategies have been useful to alter chromatographic properties, which can improve RNA separation and indirectly improve IEs by causing RNA to elute to the ESI source with higher organic solvent content. However, quantitative assessment of IE enhancement has not been reported for RNA derivatization strategies. Moreover, there are no customizable derivatization platforms that have been specifically explored to deliberately increase IEs of RNA oligonucleotides that ionize poorly.

In this study, we developed a ligation-based approach for RNA oligonucleotide labeling that increases MS sensitivity via improved IEs. In this approach, chemically modified oligodeoxynucleotides serve as MS signal enhancers upon ligation to RNA molecules. Signal enhancers were produced either by the incorporation of n-alkyl carboxylic acids in 3′ amino modified DNA oligonucleotides through amide bond formation or by the incorporation of alkene functionalized imidazoles on 3′ thiol modified oligonucleotides. We tested ethyl, hexyl, octyl, and decyl modified signal enhancers and found that decyl groups resulted in the greatest increase in ionization efficiencies for oligonucleotides, yielding an ∼15-fold increase in MS signal compared to unmodified oligonucleotides. Despite the presence of a bulky decyl modification, signal enhancers were readily appended to RNA using an RNA ligase to afford a ∼3-fold increase in MS sensitivity. Finally, we report an optimized sample preparation workflow to ligate signal enhancers to tRNA(Phe) digestion products originating from *Saccharomyces cerevisiae*, improving LC-MS/MS detection of sequence-informative digestion products and tRNA modification mapping in as low as 200 ng of tRNA input compared to a label-free method.

## Materials and Methods

### Chemicals and reagents

All reagents were used without further purification. DNA and RNA oligonucleotides were purchased from Integrated DNA Technologies (Coralville, IA, USA) with standard desalting purification. Hexanoic acid, octanoic acid, decanoic acid, and 2,2-dimethoxy-2-phenylacetophenone were purchased from TCI America (Portland, OR, USA). RNase T1, SMART digest RNase T1, N-hydroxysuccinimide long chain biotin (NHS-LC-biotin), Sulfo-N-hydroxysuccinimide (sulfo-NHS), 2-morpholin-4-ylethanesulfonic acid (MES), LC-MS grade ammonium acetate, water, acetonitrile, and Trizol reagent were purchased from Thermo Fisher Scientific (Waltham, MA, USA). 1-ethyl-3-(3-dimethlyaminopropyl)carbodiimide (EDC) was purchased from Advanced Chemtech (Louisville, KY, USA). Alkaline phosphatase was purchased from Worthington Biochemical (Lakewood, NJ, USA) and T4 RNA Ligase 1 was purchased from New England Biolabs (Ipswich, MA, USA). Streptavidin Sepharose beads were purchased from Cytiva (Marlboro, MA, USA). Transfer RNA Phenylalanine from brewer’s yeast was purchased from Millipore Sigma (Burlington, MA, USA).

### Chemical Derivatization of DNA oligonucleotides

In this study, two different derivatization methods were explored. In one approach, ion-tagged oligonucleotides were prepared by thiol-ene click reactions as described previously.^41^ Briefly, 400 nmol of allylimidazolium bromide salt and 4 nmol of reduced 3′ thiol modified DNA oligonucleotide were spotted on a 96 well plate and exposed to 365 nm light for 1 h. Reactions were diluted 10-fold with water and loaded onto 18% denaturing polyacrylamide gel at 110 V and visualized by UV shadowing.

In a second approach, 3′ alkyl amide DNA conjugates were prepared using amide coupling chemistry in 0.1 M MES buffer (pH 6). To activate the carboxylic acid, 10 µL of 40 mM acid prepared in 50/50 acetonitrile/MES buffer, 1 µL of 400 mM sulfo-N-hydroxysuccinimide (sulfo-NHS) in MES buffer, and 9 µL of 400 mM 1-ethyl-3-(3-dimethlyaminopropyl)carbodiimide (EDC) in MES buffer were combined in a microcentrifuge tube and placed on a rotary shaker for 15 min. The pH was then increased to 8 by adding 15 µL of 0.5 M NaHCO_3_. A 5 µL aliquot of 4 mM 3′ amino oligonucleotide was added to the mixture for a final volume of 40 µL and the reaction was placed on the rotary shaker overnight.

Both types of derivatized oligonucleotides were purified via reversed-phase liquid chromatography using a Zorbax Eclipse Plus C18 column (2.1 × 100 mm, 3.5 µm). A 10 min 0-20% of mobile phase B linear gradient elution was used with 10 mM ammonium acetate as mobile phase A and acetonitrile as mobile phase B, at 35 °C at a flow rate of 0.5 mL/min. Pure fractions were dried in a Quattro MiVac vacuum centrifuge concentrator and resuspended in water before analysis. Conjugates were characterized via flow injection analysis with high resolution mass spectrometry with the Agilent 6530B system described below (Figure S1).

### Nuclease Biotinylation and Ribonuclease Digestion of RNA

To allow enable facile removal of nucleases following RNA digestion and ligation, we explored two RNase T1 immobilization strategies. For one strategy, RNase T1 was biotinylated using previously established procedures.^42^ Briefly, NHS-LC-Biotin was added in 15-fold molar excess relative to RNase T1 in a solution in 50 mM HEPES buffer and incubated at room temperature for 30-90 min before quenching the reaction with 10 mM Tris HCl. Streptavidin Sepharose beads were washed three times with water before combining 100 µL of beads with 4 µL (∼400 U) of biotinylated T1. For RNA digestion, 25 µL of the bead mixture was added to the RNA of interest and incubated at 37 °C at desired time in a final volume of 100 µL of water before centrifuging at 8,000 x *g* for bead removal. The resulting supernatant was speed vacuumed to dryness before analysis.

To ensure complete removal of RNase T1, commercially available RNase T1 immobilized to magnetic particles was used.^43^ RNA was digested in microcentrifuge tubes in a final volume of 100 µL with 1-2 µL of prewashed immobilized T1 at 37 °C for 5 mins. RNase T1 was removed after digestion with a magnet placing the supernatant in a fresh microcentrifuge tube. Digestion products were speed vacuumed to dryness and resuspended in T4 RNA ligase 1 buffer. To remove residual 3′ phosphates, 50 U of T4 PNK were added and incubated at 37 °C for 1 h at a final volume of 10 µL in a PCR tube. The resulting RNA oligonucleotides were analyzed directly by LC-MS/MS or used as substrates for ligation with chemically derivatized DNA oligonucleotides.

### Ligation with T4 RNA Ligase 1

Ligation of 5′ phosphorylated DNA donors to the 3′ terminus of RNA acceptors was carried out using T4 RNA Ligase 1 (T4 RNL1) in the presence of ATP. In a PCR tube, a 20 µL reaction was prepared consisting of 15% DMSO, 1x T4 RNA Ligase Reaction Buffer, 1 mM ATP, 20-100 pmol of RNA acceptor, 100-1000 pmol of DNA donor, 1 µL of RNase Inhibitor Murine, and 2 U/µL of T4 RNL 1. Reactions were incubated for 2 h at 25 °C and then overnight at 16 °C in an Applied Biosystems Miniamp thermocycler.

### Determination of ESI ionization efficiencies for oligonucleotides

The ionization efficiencies of n-alkyl-modified DNA conjugates and their ligation products with RNAs were assessed using an Agilent 1260 Infinity II equipped with a quaternary pump (G7111B), automated vial sampler with a built-in column heater (G7129A), and a diode array detector (G7115A) in line with an Agilent 6120 single quadrupole mass spectrometer equipped with an API-ES source. Derivatized n-alkyl-modified DNA conjugates were injected via flow injection analysis at a flow rate of 50 µL/min in 40/60 10 mM ammonium acetate/ ACN. The electrospray source was operated in negative mode, with a drying gas flow rate of 12 L/min at a temperature 300 °C and a fragmentor voltage of 70 V. The nebulizer pressure was set to 35 psig. The capillary voltage was 3000 V. The quadrupole temperature was set to 100 °C. The cycle time was set for a peak width of 0.10 min with 50% of the cycle time in full scan mode from 100 – 1000 *m/z* and 50% of cycle time in selected ion monitoring for the *m/z* corresponding to the 2^nd^ and 3^rd^ charge states. Peak volumes were determined by summing selected extracted ion chromatogram (EIC) peak areas for the 2^nd^ and 3^rd^ charge states as well as sodium adducts, which were the most dominant signals for all n-alkyl-modified DNA conjugates.

### Modification mapping of yeast tRNA(Phe)

For analysis of tRNA, the same LC system described above was coupled to an Agilent 6530B QTOF equipped with a Dual AJS ESI source in negative mode. Digestion products were separated on a Phenomenex (Torrance, CA, USA) Biozen Oligo Column (2.1 × 100 mm, 2.6 µm) using 10 mM ammonium acetate as mobile phase A and a premixed mobile phase B consisting of 50% mobile phase A and 50% acetonitrile. A linear gradient elution program consisting of a 5 min hold at 0% B followed by linear ramps of 0-20% in 15 min, 20-30% B in 5 min, 30-55% B in 35 minutes at 0.25 mL/min at 60 °C. The LC eluent was diverted to waste for the first 5 min. The column was washed with 80% acetonitrile for 2 min and then reequilibrated with mobile phase A for 20 minutes. The nebulizer gas was set at 35 psi. The drying gas flow rate was 12 L/min at a temperature of 300 °C. The sheath gas flow rate was 12 L/min at a temperature of 350 °C. A nozzle voltage of 2000 V was applied followed by a capillary voltage of 3000 V, a fragmentor voltage of 100 V, a skimmer voltage of 65 V and an octopole voltage of 750 Vpp. Mass spectra were acquired in data dependent acquisition mode with an overall duty cycle consisting of a 0.33 s MS1 scan (*m/z* 400-3200) followed by 2-4 1 s MS2 scans (*m/z* 100-3200). A 4 *m/z* isolation window was used with CID fragmentation using optimal collision energies previously reported for RNA on the same model of MS instrument.^44^ MassHunter Qualitative Analysis 10.0 or Proteowizard MSConvert were used to convert MS data into .mgf file format. Peak areas were calculated by extracting chromatograms for ions in most abundant charge states corresponding to the monoisotopic and major isotopes with a 50 ppm window in MassHunter Qualitative Analysis 10.0.

With minor modifications, the open-source software package, Pytheas,^44^ was used to generate *in silico* digestions of tRNA and compare predicted MS2 spectra with data collected. For the analysis of ligation products, we added a custom python script to the *in silico* digestion workflow that creates additional search queries for ligation products based on the expected digestion products as well as a user specified donor oligonucleotide sequence. Python script and additional data visualization R scripts for generation of graphics are publicly accessible (https://github.com/mshari03/RNA_MS_signal_enhancement). The tRNA digestion and ligation mass spectrometry data sets were deposited in mgf format on the ProteomeXchange consortium via the PRIDE partner repository with the dataset identifier PXD076675.

## Results and Discussion

### Optimizing the structure of signal-enhancing oligonucleotides for improved ionization efficiency

RNA oligonucleotides have a high capacity for hydrogen bonding and polyanionic backbones that contribute to poor ESI-MS sensitivity and ultimately require the analysis of large quantities of RNA sample. We aimed to overcome this challenge by adding a hydrophobic functional group to the RNA structure, thereby increasing its surface activity and ionization efficiency. Owing to the large number of exocyclic amines and 2′ OH groups present on RNA, there are a limited number of direct conjugation strategies that are both site selective and result in high yields.^45^ Although new approaches are being developed for direct derivatization of endogenous RNA,^46–48^ we reasoned that post-synthetic derivatization of amino- or thiol-modified oligonucleotides would provide a highly customizable derivatization strategy and enable access to diverse functional groups for investigating their impact on IE. To identify which chemical functional groups have the largest impact on ionization efficiencies, we first coupled a 5-mer DNA oligonucleotide bearing a 3′ amino handle to a series of n-alkyl carboxylic acids two, six, eight, and ten carbons in length (Figure 1A). Carboxylic acids with more than ten carbons were only sparingly soluble in the primarily aqueous reaction conditions, so their synthesis was not explored further. The ionization efficiencies of the n-alkyl-derivatized DNA oligonucleotides were studied using flow injection ESI-MS under isocratic conditions to control for the impact of solvent composition on ESI. When compared to an unmodified DNA oligonucleotide, the incorporation of alkyl chains into the DNA structure led to an increase in MS signal intensity that generally correlated with length of the alkyl chain (Figure 1B). The correlation between chain length and ESI-MS response provides further evidence that increased surface activity mediated by the analyte’s amphiphilic nature contributes to increased ionization efficiency. The decyl conjugate resulted in a 14.9-fold increase in sensitivity as compared to DNA oligonucleotide with an unreacted amino handle as measured by the slope of calibration curves. We also used thiol-ene chemistry to couple 3′ thiolated oligonucleotides with allylimidazolium salts bearing various length alkyl moieties at the N3 position,^41^ as we were interested in exploring whether the addition of the positively charged imidazolium group would further improve IE. While the alkyl imidazolium constructs also showed increasing IEs with alkyl chain length, similar IEs were observed for the imidazolium derivatives in both positive and negative mode, indicating little benefit to adding the imidazolium group (Figure S2). In addition, thiol-ene reactions proved difficult to scale up and their purification from oxidation byproducts discouraged further investigation of these conjugates. We thus selected the decyl-functionalized DNA oligonucleotide as a signal enhancer for further experiments since it conferred the largest increase in IE, was readily synthesized using carbodiimide coupling, and purified at a relatively large scale (tens of nanomoles).

**Figure 1.**
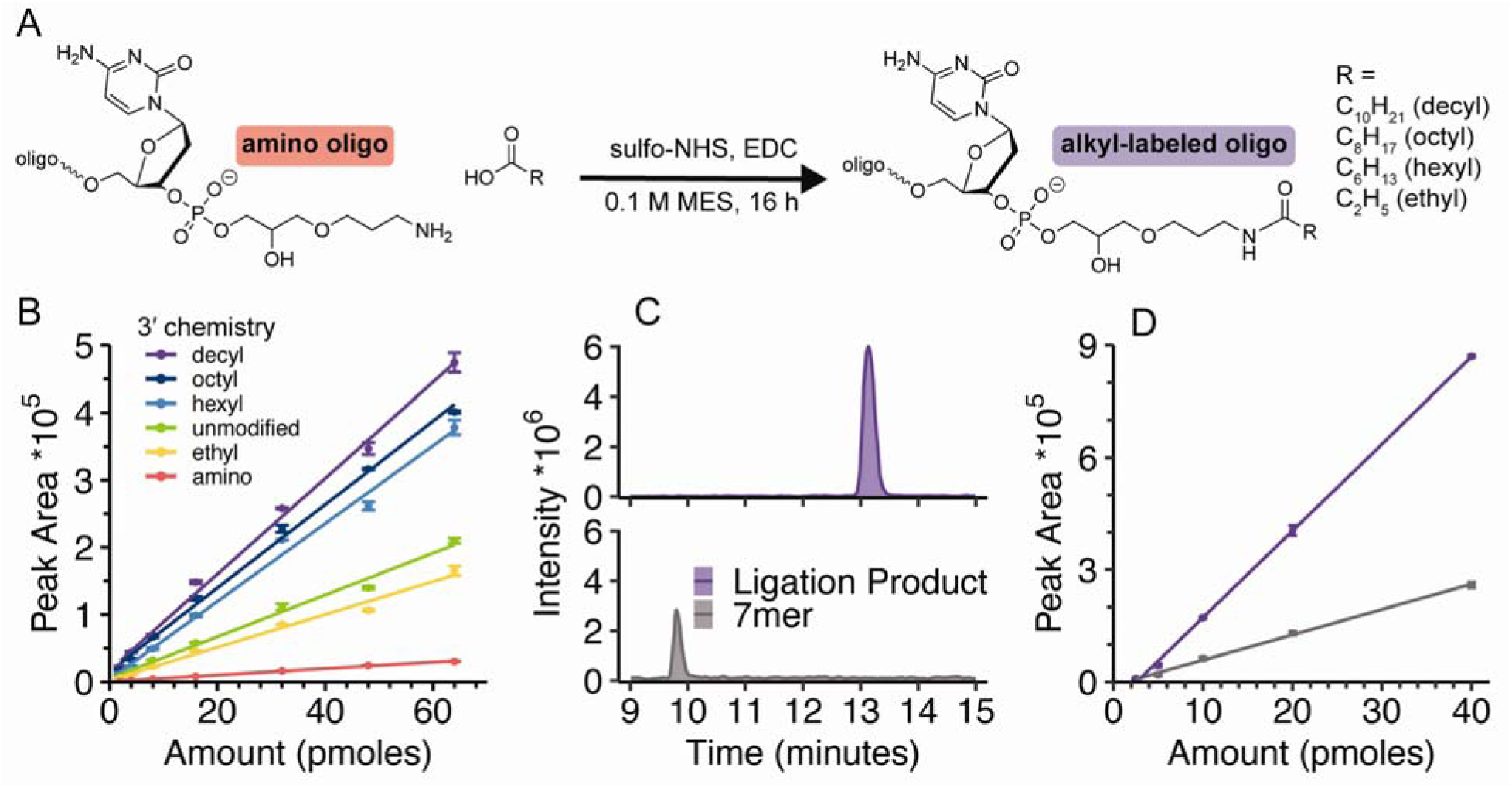
Derivatization platform to increase ionization efficiency of RNA oligonucleotides. (A) Schematic of derivatization of 3′ amino DNA oligonucleotides with carboxylic acids. (B) Calibration curves for ESI response of various signal enhancing oligonucleotide derivatives using flow injection analysis. (C) Example EICs of unmodified RNA acceptor and decyl modified ligation product. For both the 7mer RNA acceptor and the ligation product, EICs corresponding to 2 most abundant charge states as well as potassium adducts were generated using a 50 ppm window. (D) Calibration curves for LC-MS response of the modified ligation product and the corresponding RNA standard.

### Ligation with signal-enhancing oligonucleotides increases MS sensitivity for RNA

To make our signal enhancement approach broadly applicable to RNA samples, we pursued an enzymatic ligation approach to functionalize any RNA bearing a 3′ OH group. Although there is extensive work in optimizing adapter ligation for NGS approaches, this strategy is less commonly used in sample preparation for LC-MS. We thus selected T4 RNA Ligase 1 for our workflow as it is well suited for ligating single-stranded RNA acceptors of interest with RNA or DNA donors for signal enhancement. Reaction time and amount of ligase were optimized by monitoring the kinetics of ligation to an RNA standard: the majority of RNA acceptor was consumed within 2 h when 2 U/µL of ligase was used (Figure S3). Additionally, the ratio of donor to acceptor had a strong impact on ligation efficiency (Figure S4). The decyl modified signal enhancer was readily and quantitatively ligated to an RNA standard despite the presence of the bulky 3′ alkyl modification when ratios of 2 to 1 or greater were used (Figure S4). To determine whether IEs for RNA could be improved by attaching a signal-enhancing oligonucleotide, we compared the signal intensity from an unmodified 7-mer RNA standard to its ligation product. Ligation of the decyl-modified signal enhancer resulted in increased MS signal as well as increased retention under reversed phase LC conditions without ion pairing reagents (Figure 1C). In general, peak areas reported were calculated by summing EIC peak areas for the two most abundant mass to charge ratios (Table S1). Calibration curves showed a 3.4-fold increased MS sensitivity for the modified ligation product compared to the unmodified 7-mer RNA (Figure 1D). Additionally, the MS2 spectra of the ligation products confirmed ligation of the signal enhancer to the RNA standard and illustrated that our labeling strategy did not interfere with MS-based sequencing of the target RNA (Figure S5A). Thus, ligating RNA targets with signal enhancers increases LC retention, improves MS signal intensity for RNA, and permits CID fragmentation for MS2 that is necessary for RNA sequencing.

Encouraged by these results, we investigated the MS signal intensities of a mixture of RNA standards pre- and post-ligation to determine the applicability of the method to several RNA species at once, as well as the impact of the signal enhancer oligonucleotide on RNA ionization efficiency. We chose to investigate 7-mer, 9-mer, and 13-mer RNA species which are representative of typical RNA fragment lengths produced by RNase T1 digestion. Again, the ligation of the signal enhancing oligonucleotide improved signal and increased retention for all three RNA oligonucleotides in the standard mixture (Figure 2B). Additionally, after ligation with the signal enhancer, no remaining unlabeled RNA acceptor was observed by PAGE or LC-MS, indicating highly efficient ligation (Figure 2A and Figure S6). Ligation to the signal enhancer resulted in a ∼2-fold increase in peak volume for all three RNA analytes tested (Figure 2C), suggesting that the signal enhancement conferred does not depend on the length of the RNA acceptor within the studied size range. Interestingly, we observed a shift in the charge state distribution after ligation, which can be explained by lengthening the RNA target by an additional five phosphate groups from the signal enhancer structure. For example, the 13-mer RNA, initially observed predominantly in the −3 charge state, was primarily detected in its −4 charge state after ligation (Figure 2D). Overall, these results demonstrate the applicability of signal enhancement to a single sample containing a mixture of several RNAs of different lengths.

**Figure 2.**
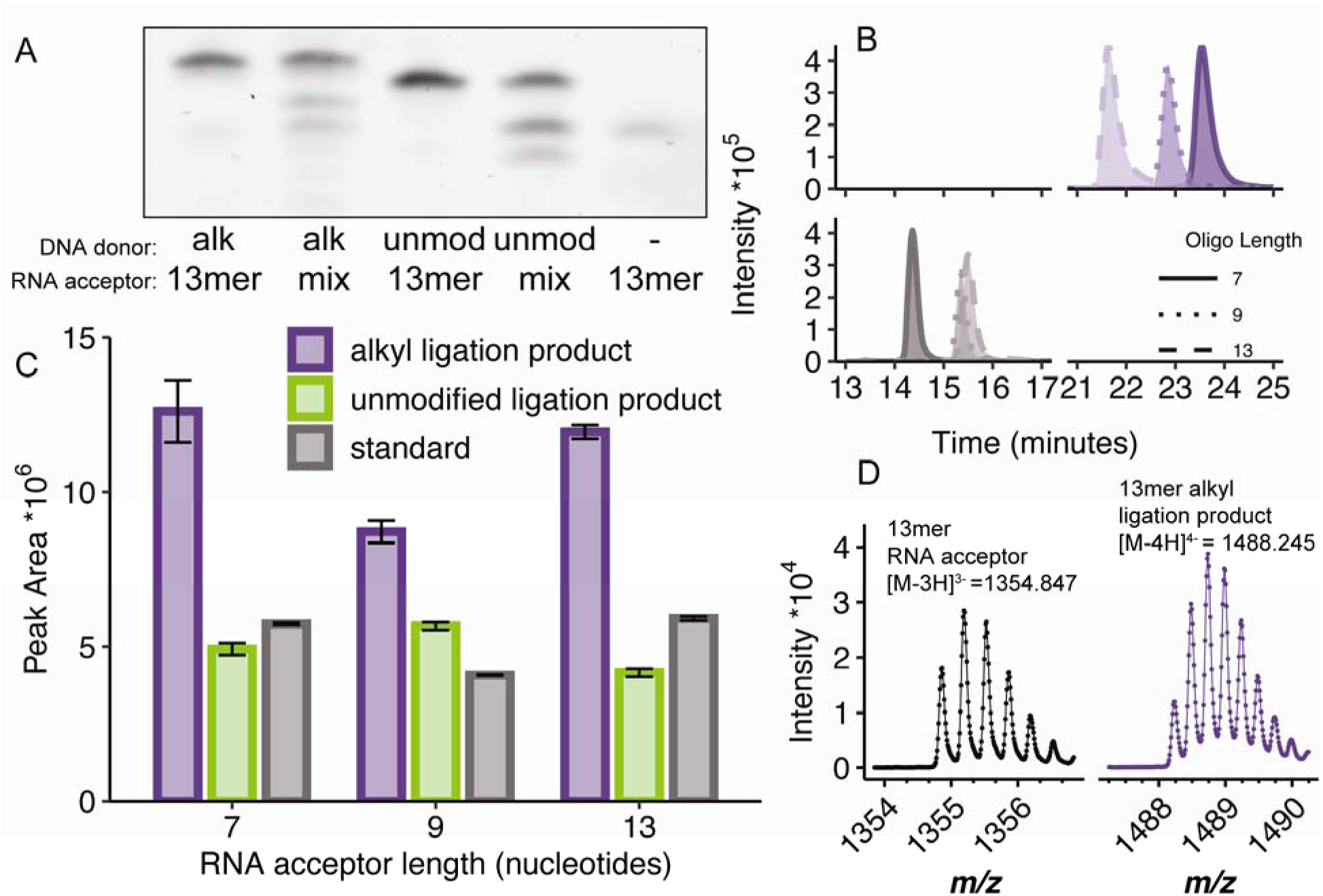
Impact of derivatization on a mixture of RNA standards. (A) Ligation reaction completion was confirmed by PAGE with SYBR gold staining. (B) RNA standards before and after ligation were separated via reversed phase LC prior to ESI in the absence of ion pairing agents. EICs corresponding to the 2 most abundant charge states for both the monoisotopic mass and the M+1 isotope were generated with a 50 ppm window. (C) Change in ESI response after ligation with either an unmodified 5 nucleotide DNA donor or a decyl modified DNA donor (D) Example mass spectra of unmodified RNA acceptor compared to the decyl modified ligation product.

### Optimization of sample preparation conditions for enhancing signals of RNA digestion products

We were motivated to incorporate ligation-based signal enhancement to popular LC-MS/MS workflows for modification mapping of endogenous RNAs which involve sequence-selective digestion with nucleases. However, when directly applying RNA digests to ligation with signal enhancers, we did not observe ligation. We hypothesized that RNase T1 remained active in solution, so we explored strategies to immobilize ribonuclease T1 to enable facile removal prior to ligation.^43,49,50^ A cost effective method for immobilization involves biotinylation of RNase T1 and removal with streptavidin coated Sepharose beads (Figure S7A-B). Nuclease immobilization has the previously reported advantage^43,50^ of generating diverse digestion products via missed cleavages (Figure S7C-D). Limited ligation efficiencies were observed with biotinylated RNase T1, which could be caused by residual free T1 cleaving ligation products. Thus, to ensure complete removal of RNase T1 from the sample, we turned to commercially available functionalized magnetic beads that have been previously used for LC-MS analysis of longer RNAs.^43^

For complete ligation, T4 RNA Ligase 1 requires that RNA acceptors contain a 3′ alcohol for phosphodiester bond formation with a 5′ phosphate donor. Popular nucleases like RNase T1 and RNase A cleave 3′ to specific nucleotides creating a mixture of linear phosphates, cyclic phosphates and free 3′ alcohols. There is evidence that partial digestion by immobilized nucleases results in a higher proportion of cyclic and linear phosphates compared to free alcohols.^50^ Bacterial alkaline phosphatase has been used previously to remove phosphates prior to LC-MS of digested RNA,^51^ but this enzyme is challenging to inactivate and will remove 5′ phosphates from the donor molecule preventing ligation. However, we found that T4 Polynucleotide Kinase (T4 PNK) effectively removes 3′ phosphates (Figure S8) and is more readily heat inactivated. The optimized workflow (Figure 3A) was applied to a 31 nucleotide RNA standard to enable high efficiency ligation (Figure 3B). Ultimately, multiple enzymatic transformations of the RNA of interest permitted high efficiency ligation with a customizable signal enhancing DNA oligonucleotide.

**Figure 3.**
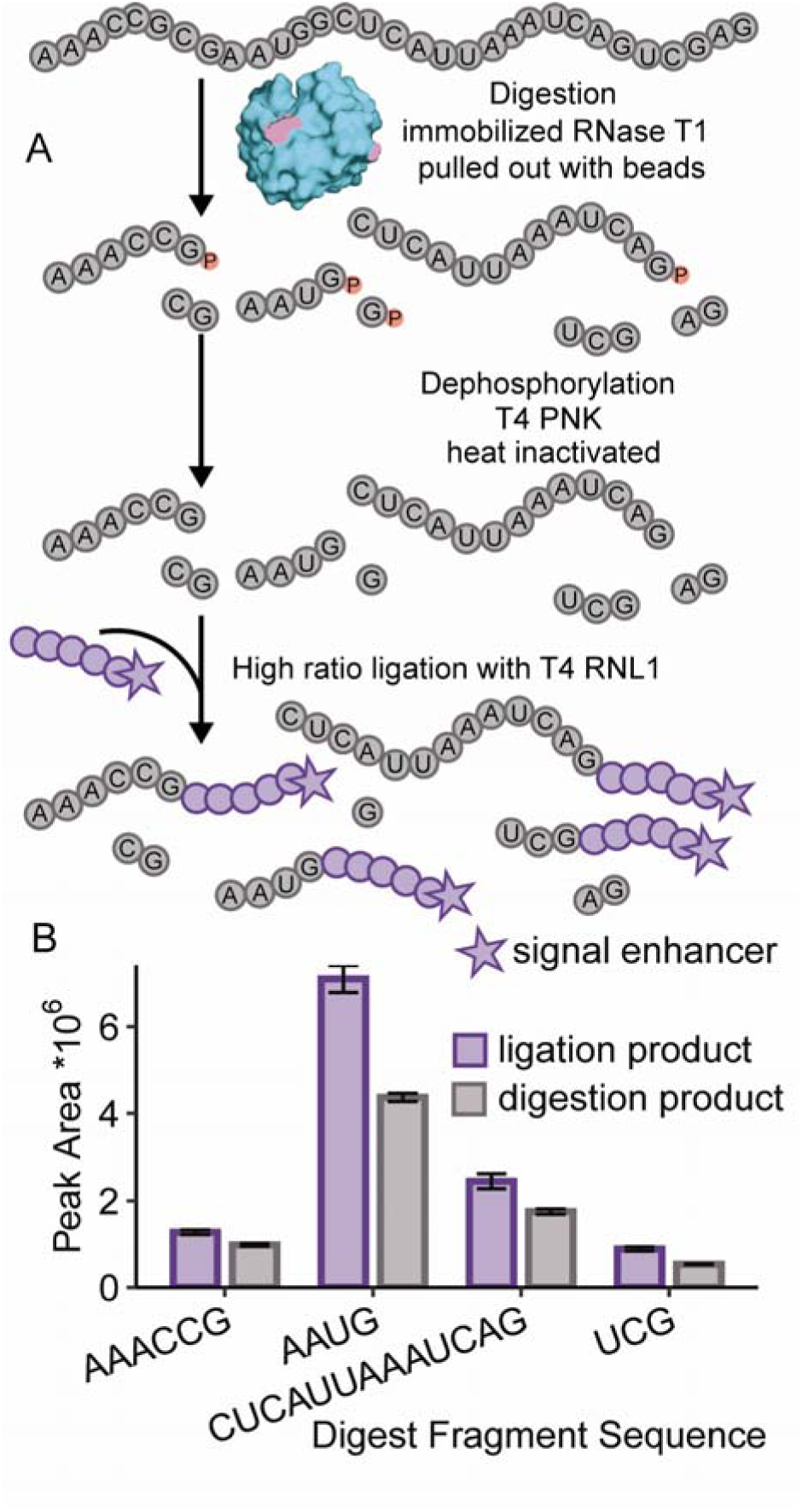
Sample preparation strategy for signal enhancement for bottom-up RNA analysis by MS. (A) Schematic of sample preparation steps. Structure of ribonuclease T1 shown with available lysine residues for conjugation highlighted in pink. (B) When applied to a 31mer RNA standard, the resulting EIC peak areas were compared to RNase T1 digestion products that did not undergo ligation. The average of three technical replicates is shown, error bars represent ±1 SD. Peak areas were calculated from EICs for the two most abundant charge states for each oligonucleotide species.

### Application of ligation-based signal enhancement to endogenous modified tRNA

We next applied our MS signal enhancement workflow to a tRNA harboring multiple post-transcriptional modifications. tRNAs are highly modified adaptor molecules that convert genetic information to amino acids. Modifications of tRNAs are dynamically altered in response to stress and have been implicated in a variety of human diseases,^52^ increasing the demand for methods capable of sensitive mapping of tRNA sequences inclusive of modifications. We thus applied our ligation-based signal enhancing strategy to the 76 nucleotide tRNA(Phe) from *S. cerevisiae*, which is known to contain 13 modifications.^53^ Ribonuclease T1 creates 21 expected digestion products (Figure 4A), and without signal enhancement, we observed all tRNA digestion products larger than 2 nucleotides using 500 ng of tRNA starting material. After ligation with signal enhancing oligonucleotides, we observed ligation products for all modified tRNA fragments larger than 3 nucleotides. Furthermore, a 1.3 to 8 fold change increase in peak volume was observed for ligation products as compared to corresponding T1 digest fragments analyzed by label-free LC-MS/MS (Figure 4B). When only 200 ng of starting material is used, increased signal is observed for all tRNA digestion products except for the fragment corresponding to the anticodon stem loop, which was not detected by either method at that input amount (Figure S9, Table S1). Additionally, at this lower amount, 5 out of the 6 detected RNA ligation products had matching MS2 spectra. Since many tRNA species are low abundant or hypomodified in limited samples, a signal enhancing approach is necessary for detection and thorough modification analysis.

**Figure 4.**
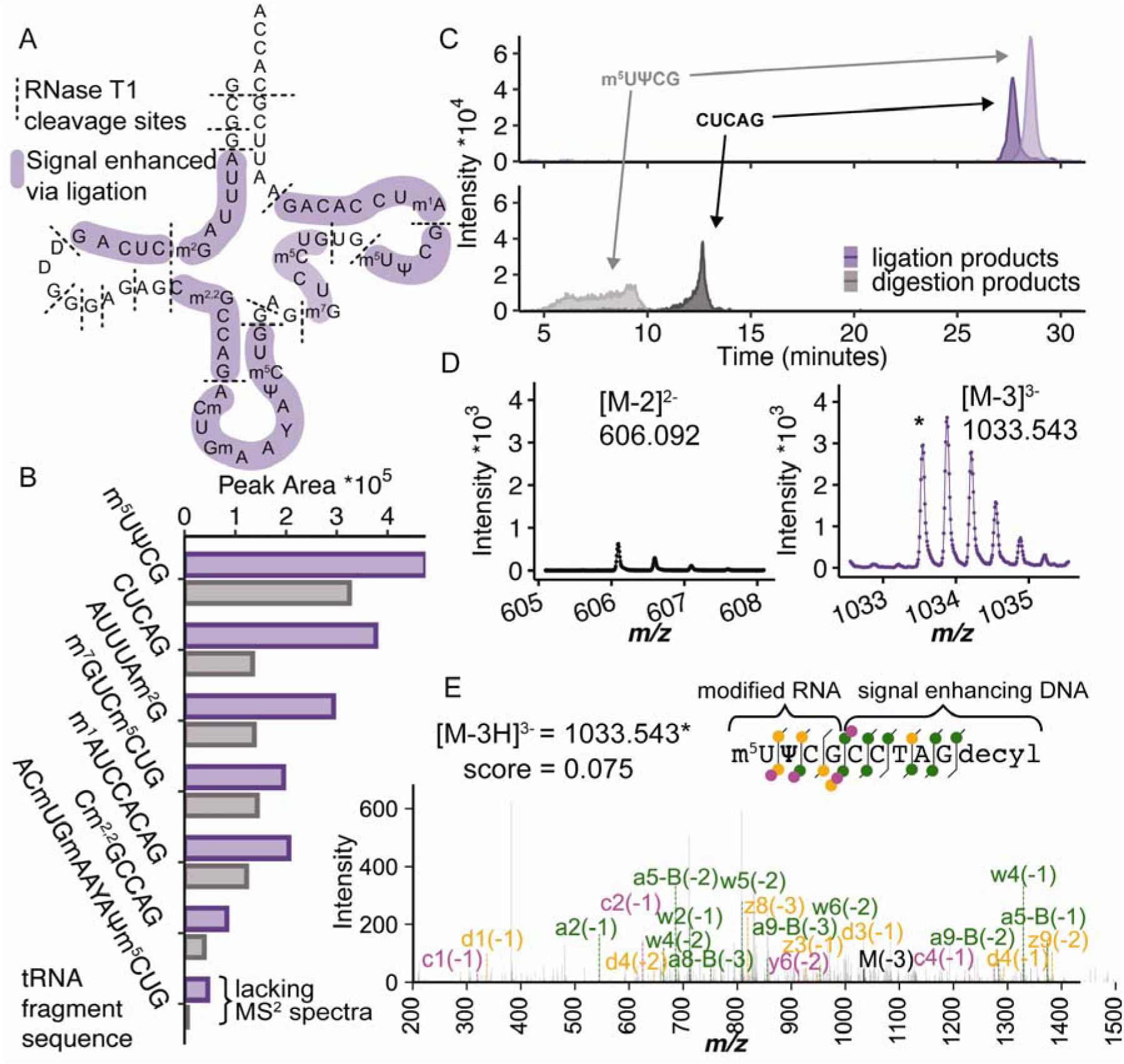
Application of the ligation-based signal enhancement strategy for LC-MS/MS analysis of tRNA(Phe) (A) Schematic of tRNA(Phe) from *S. cerevisiae* with known modifications shown as abbreviations. Expected RNase T1 cleavage sites are shown as dashed lines and fragments observed are highlighted. (B) EIC peak areas for the two most abundant charge states of RNA oligonucleotides before and after ligation of signal enhancing DNA oligonucleotide (Ψ signifies pseudouridine, Y signifies wybutosine). (C) Example EICs for ligation products (top) and RNase T1 digestion products without ligation (bottom). (D) Example mass spectra for [m^5^U]ΨCG before and after ligation. (E) Example MS2 spectra confirming sequence identity of ligation product.

We found that ligation products of [m^5^U]ΨCG and CUCAG displayed increased LC retention and improved peak shapes compared to analysis without ligation (Figure 4C). These results show that in addition to signal enhancement, our ligation-based derivatization strategy has an advantage of improving chromatographic properties of RNA molecules in the absence of ion pairing agents. The identities of these ligation products were confirmed with high mass accuracy and expected isotopic distributions (Figure 4D). Moreover, we further confirmed 6 ligation products via MS2 spectral matching with high coverage of sequence ladder fragments (Figure 4E and Figure S10). Primarily a- and w-ions were observed in the DNA signal enhancer portion whereas more c- and y-ions were observed in the RNA portion due to the known propensity of DNA base loss.^54^ Since many spectral scoring algorithms are not designed for these chimeric sequences, the difference in fragmentation pattern likely leads to lower spectral scores. Nonetheless, ligation of signal enhancers to a tRNA digest demonstrates the versatility of this sample preparation approach to enhance ionization efficiency as well as increased reverse-phase chromatographic retention of modified RNA molecules.

### Conclusions

RNA sequencing by MS is inherently limited by poor RNA ionization efficiency and the need for ion pairing agents that often contaminate LC-MS instrumentation. These limitations have created a demand for methods that improve MS sensitivity and LC separations of RNA that enable analysis of low abundance RNA samples. In this study, we developed a ligation-based platform to improve LC-MS/MS sequencing of RNA that capitalizes on chemically customizable oligodeoxynucleotide signal enhancers. We investigated the impact of a series of functional groups on the ionization efficiencies of short oligodeoxynucleotides and found that the addition of hydrophobic groups can greatly improve ionization. Signal enhancers functionalized with decyl groups showed the best performance providing a 14.9-fold increase in sensitivity compared to unmodified DNA. When signal enhancers were ligated to RNA standards, sensitivity gains of 2-4 fold were observed. An additional advantage of the ligation-based approach was improvements in LC retention and chromatographic peak shape in the absence of ion pairing reagents. Thus, we anticipate this strategy will be useful for general-purpose labs that do not have access to dedicated LC-MS instruments and want to avoid LC methods involving ion pairing reagents. Furthermore, when to a tRNA from *S. cerevisiae*, our method provided signal enhancement for numerous sequence-informative fragments bearing post-transcriptional modifications. More sensitive measurements will allow more accurate quantification of RNA modification stoichiometries which are crucial to discern their function. A major advantage of our workflow over existing RNA derivatization methods is the customizability of the functional groups on the signal enhancer. Chemical groups can be tailored for specific purposes including improved ionization and retention, or perhaps RNA fragmentation or other objectives needed to enhance LC-MS based methods for epitranscriptomic measurements.

## Supporting Information

High resolution mass spectra of conjugates; ionization efficiency of ion tagged oligonucleotides; optimization of enzymatic sample preparation; annotated tandem mass spectra of ligation products.

## Author Contributions

The manuscript was written through contributions of all authors, who approved the final version of the manuscript.

## Acknowledgments

K.D.C. acknowledges the Arnold and Mabel Beckman Foundation for support through the Beckman Young Investigator Award Program.

---

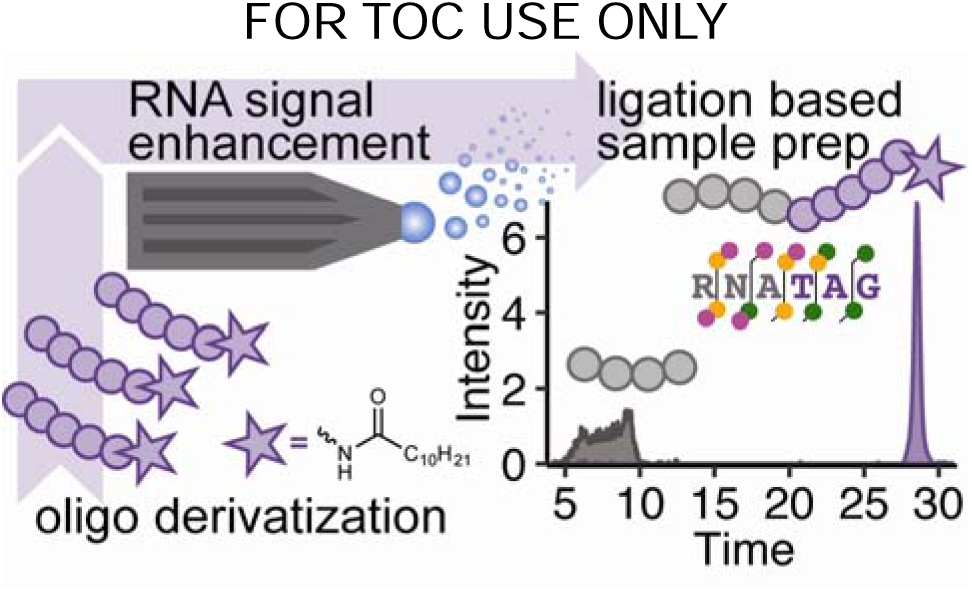

## Notes

https://github.com/mshari03/RNA_MS_signal_enhancement

### Competing Interest Statement

The authors have declared no competing interest.

