## Supporting Information for "Enzymatic Ligation Strategy to Enhance Electrospray Ionization Efficiency and Liquid Chromatography-Mass Spectrometry of DNA and RNA Oligonucleotides"

Authors and Affiliations

Max D. Sharin, Nina J. Fitzgerald, Stephen M. Kennedy, Isabella G. Park, Kevin D. Clark*

Department of Chemistry, Tufts University, Medford, MA 02155, United States

**Table of Contents**

(p. 2) **Protocol** for Polyacrylamide Gel Electrophoresis

(p. 2) **Protocol:** Isolation of 23S rRNA from *E. coli*

(p. 4-6) **Table S1:** Oligonucleotide sequences and mass to charge ratios used to generate extracted ion chromatograms

(p. 7) **Figure S1:** High resolution mass spectra of amino alkyl conjugates

(p. 8) **Figure S2:** Synthesis of imidazolium tagged oligonucleotides and ionization efficiency measurements

(p. 9) **Figure S3:** Ligation reaction kinetics monitored by LC-MS

(p. 10) **Figure S4:** Polyacrylamide gel electrophoresis of ligation of signal enhancer to an RNA standard

(p. 11) **Figure S5:** Tandem mass spectra for ligation products of RNA standards

(p. 12) **Figure S6:** Extracted ion chromatograms for RNA acceptors remaining after ligation

(p. 13) **Figure S7:** Biotinylation of ribonuclease T1and limited digestion of *E. coli* 23S rRNA

(p. 14) **Figure S8:** Optimization of amount of T4 Polynucleotide Kinase needed to remove 3′ phosphates

(p. 15) **Figure S9:** Tandem mass spectra of ligation products derived from tRNA

**Figure S10:** Peak area comparisons between 500 and 200 ng of tRNA inputs

**Protocol:** for polyacrylamide gel electrophoresis

Polyacrylamide gels (18%) were cast in house by mixing 4.8 g urea, 4.5 mL of 10X Tris-Borate-EDTA buffer, 4.5 mL of 40% acrylamide/bis solution 29:1. After degassing the mixture in a sonicator, 100 µL of 10% ammonium persulfate in water and 10 µL of N,N,N,N-tetramethylethylenediamine (TEMED) were added to initiate polymerization before loading a Biorad mini protean gel cast (8 cm x 10 cm) with 1.0 mm spacer plates to set overnight. Samples were premixed with an equal volume of 8M urea and loaded on after prerunning for 5-10 minutes. Gels were run for 75 minutes at 110 V on a Biorad mini-Protean tetra cell system, stained with SYBR gold for 5 minutes, and imaged with an invitrogen iBright 1500 gel imager.

**Bacterial culture and rRNA isolation**

NEB 5α competent *Escherichia coli* were purchased from New England Biolabs (Ipswich, MA, USA). Bacteria were transformed with pUC19 according to the manufacturer’s instructions to confer ampicillin resistance. Single colonies were picked from LB agar plates with ampicillin and grown up in LB Broth with 50 µg/mL ampicillin at 37 ˚C with shaking at 250-275 rpm until exponential growth phase with an OD600 of 0.4 - 0.7. Cells were pelleted at 5,000*g* for 10 minutes at 4 ˚C. Cell lysis was performed using Bacterial Protein Extraction Reagent in buffer A (20 mM Tris pH 7.5, 300 mM NH_4_Cl, 10mM MgCl_2_, 0.5 mM EDTA, 6 mM β-mercaptoethanol) via two rounds of freeze thawing on dry ice and ice. rRNA was isolated using sucrose gradient ultracentrifugation following established protocols.^1^ Briefly, after clarifying the cell lysate via centrifugation at 21,300*g*, crude ribosomes were pelleted at 100,000*g* for 1 h at 4 ˚C. For subunit purification, crude ribosomes were resuspended in dissociating buffer A (containing 1 mM MgCl_2_) and loaded on top of a 10-40% (w/v) sucrose gradient prepared in buffer A^2^ and centrifuged at 88,000*g* for 16 h at 4 ˚C. After needle drop fractionation, rRNA was purified using the Trizol method. RNA pellets were redissolved in water, the concentration was determined via A260 measurement, and when necessary, rRNA was concentrated using Amicon Ultra 10 kDa MWCO filters (Millipore). rRNA was aliquoted and placed at -80 ˚C for storage and thawed on ice before use.

Table S1: Oligonucleotide sequences and mass to charge ratios used for EIC generation. All extracted ion chromatograms were generated using Agilent MassHunter Qualitative Analysis 10.0, with a 50 ppm window by merging multiple masses into one chromatogram.

| **Figure** | **identity** | **Sequence** | **Monoisotopic Mass** | **Charge States Observed** | **m/z for EICs** | **Adducts included in peak volumes** |
| --- | --- | --- | --- | --- | --- | --- |
| 1B | DNA Conjugate | dAdGdTdCdC | 1462.291 | -2 | 730.15 | +Na |
| 1B | DNA Conjugate | dAdGdTdCdC-amino | 1673.3541 | -3, -2 | 835.67, 556.78 | +Na |
| 1B | DNA Conjugate | dAdGdTdCdC-ethyl | 1715.3647 | -3, -2 | 856.67, 570.78 | +Na |
| 1B | DNA Conjugate | dAdGdTdCdC-hexyl | 1771.4273 | -3, -2 | 884.71, 589.47 | +Na |
| 1B | DNA Conjugate | dAdGdTdCdC-octyl | 1799.4586 | -3, -2 | 898.72, 598.81 | +Na |
| 1B | DNA Conjugate | dAdGdTdCdC-decyl | 1827.4899 | -3, -2 | 912.74, 608.15 | +Na |
| 1C, 1D | RNA acceptor | UCAUAUG | 2164.311 | -3, -2 | 720.429, 1081.148 | +K, +2K, +3K |
| 1C, 1D | Ligation Product | UCAUAUG-dAdGdTdCdC-decyl | 4053.7595 | -4, -3 | 1012.432, 1012.683, 1012.933, 1350.245, 1350.495, 1350.745 | +K, +2K, +3K |
| 2B, C | RNA acceptor | UCAUAUG | 2164.311 | -3, -2 | 720.429, 720.762, 1081.148, 1081.648 | None |
| 2B, C | RNA acceptor | CAAUUCUAG | 2798.405 | -3, -2 | 932.129, 932.797, 1398.194, 1398.694 | None |
| 2B, C | RNA acceptor | CUCAUUAAAUCAG | 4067.575 | -4, -3 | 1015.887, 1016.138, 1354.850, 1355.183 | None |
| 2B, C | Unmodified Ligation Product | UCAUAUG-dAdGdTdCdC | 3688.563 | -4, -3 | 921.133, 921.384, 1228.513, 1228.846 | None |
| 2B, C | Unmodified Ligation Product | CAAUUCUAG-dAdGdTdCdC | 4322.657 | -4, -3 | 1079.656, 1079.907, 1439.878, 1440.211 | None |
| 2B, C | Unmodified Ligation Product | CUCAUUAAAUCAG-dAdGdTdCdC | 5591.827 | -5, -4 | 1117.358, 1117.558, 1117.759, 1396.949, 1397.1987, 1397.4487 | None |
| 2B, C | Modified Ligation Product | UCAUAUG-dAdGdTdCdC-decyl | 4053.760 | -4, -3 | 1012.432, 1012.683, 1012.933, 1350.245, 1350.495, 1350.745 | None |
| 2B, C | Modified Ligation Product | CAAUUCUAG-dAdGdTdCdC-decyl | 4687.854 | -4, -3 | 1170.955, 1171.206, 1171.457, 1561.610, 1561.943, 1562.276 | None |
| 2B, C | Modified Ligation Product | CUCAUUAAAUCAG-dAdGdTdCdC-decyl | 5957.024 | -5, -4 | 1190.397, 1190.598, 1190.798, 1488.248, 1488.498, 1488.748 | None |
| 3B | RNA acceptor | AAACCG | 1880.330 | -3, -2 | 939.157, 939.657, 625.769, 626.102 | None |
| 3B | RNA acceptor | AAUG | 1247.220 | -2 | 622.602, 623.102 | None |
| 3B | RNA acceptor | CUCAUUAAAAUCAG | 4067.575 | -4, -3 | 1015.887, 1016.138, 1354.850, 1355.183 | None |
| 3B | RNA acceptor | UCG | 894.570 | -2 | 446.277 | None |
| 3B | Modified Ligation Product | AAACCGdAdGdTdCdC-decyl | 3769.779 | -4, -3 | 941.437, 941.687, 1255.585, 1255.918 | None |
| 3B | Modified Ligation Product | AAUGdAdGdTdCdC-decyl | 3136.669 | -4, -3 | 1044.548, 1044.881, 783.159, 783.409 | None |
| 3B | Modified Ligation Product | CUCAUUAAAAUCAGdAdGdTdCdC-decyl | 5957.024 | -5, -4 | 1190.397, 1190.598, 1488.248, 1488.498 | None |
| 3B | Modified Ligation Product | UCGdAdGdTdCdC-decyl | 2784.019 | -3, -2 | 926.998, 927.498, 694.997, 695.330 | None |
| 4B | RNA acceptor | m5UPCG | 1216.2228 | -2 | 606.092, 606.592 | None |
| 4B | RNA acceptor | AUUUAm2G | 1875.3108 | -3, -2 | 623.421, 623.754, 935.636, 936.136 | None |
| 4B | RNA acceptor | m7GUCm5CUG | 1880.3268 | -3, -2 | 625.093, 625.4263, 938.144, 938.644 | None |
| 4B | RNA acceptor | Cm2,2GCCAG | 1902.3708 | -3, -2 | 632.441, 632.7743, 949.166, 949.666 | None |
| 4B | RNA acceptor | CUCAG | 1530.2748 | -2 | 763.118, 763.618 | None |
| 4B | RNA acceptor | m1AUCCACAG | 2507.4368 | -3, -2 | 834.13, 834.463, 1251.699, 1252.199 | None |
| 4B | RNA acceptor | ACmUGmAAYAPm5CUG | 3105.6713 | -4, -3 | 1360.56, 1360.893, 1020.178, 1020.678 | None |
| 4B | Ligation Product | m5UPCG-dCdCdTdAdG-decyl | 3764.7593 | -4, -3 | 775.155, 775.405, 1033.876, 1034.209 | None |
| 4B | Ligation Product | AUUUAm2G-dCdCdTdAdG-decyl | 3769.7753 | -4, -3 | 939.928, 940.178, 1253.573, 1253.906 | None |
| 4B | Ligation Product | m7GUCm5CUG-dCdCdTdAdG-decyl | 3791.8193 | -4, -3 | 941.181, 941.431, 1255.245, 1255.578 | None |
| 4B | Ligation Product | Cm2,2GCCAG-dCdCdTdAdG-decyl | 3419.7233 | -3 | 1262.592, 1262.925 | None |
| 4B | Ligation Product | CUCAG-dCdCdTdAdG-decyl | 4396.8853 | -4, -3 | 853.669, 853.919, 1138.561, 1138.894 | None |
| 4B | Ligation Product | m1AUCCACAG-dCdCdTdAdG-decyl | 4995.1198 | -4, -3 | 1097.959, 1098.209, 1464.281, 1464.614 | None |
| 4B | Ligation Product | ACmUGmAAYAPm5CUG-dCdCdTdAdG-decyl | 5654.2078 | -5, -4 | 1194.023, 1194.223, 1492.781, 1493.031 | None |

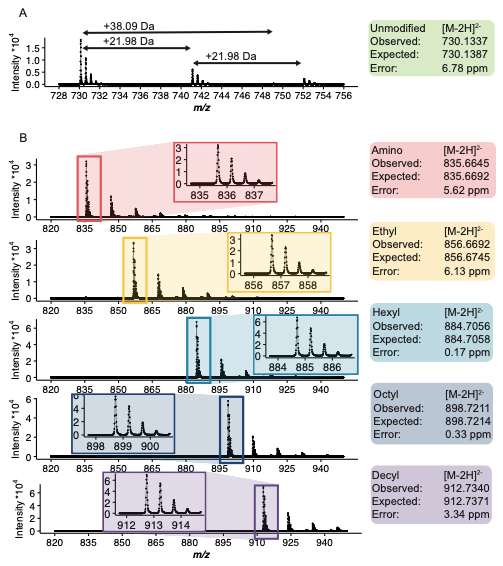
**Supplementary Figure 1.** High resolution mass spectrometry characterization of DNA oligonucleotide conjugates (A) Mass spectrum of unmodified 5 nucleotide (dAdGdTdCdC) DNA oligonucleotide with 5’ amino group (B) Mass spectra of purified n-alkyl DNA oligonucleotide conjugates, mass accuracy shown to the right.

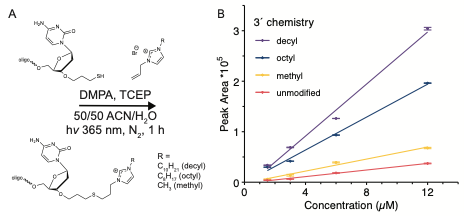
Supplementary Figure 2. Derivatization of thiol modified oligonucleotides with allyl imidazolium salts. (A) Scheme for thiol-ene reaction promoted by radical based photoinitiator (B) LC-MS calibration curves for resulting ion tagged oligonucleotides as measured by extracted ion chromatograms of the 2^nd^ charge state.

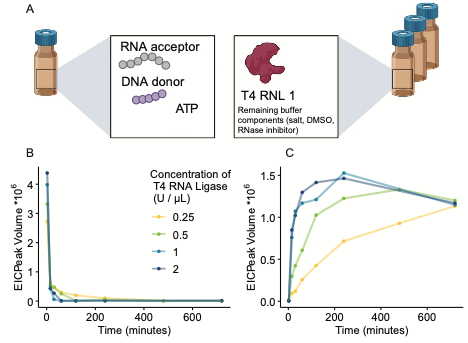

**Supplementary Figure 3:** Impact of ligase concentration on reaction kinetics. (A) Schematic of autosampler vials used in sampler injection program for kinetics analysis. One vial containing reaction components and a second vial containing appropriate concentration of ligase were mixed with the liquid handler on the HPLC and injected at set time points after mixing. (B) EIC peak volumes of RNA acceptor remaining over time. (C) EIC peak volumes of chimeric RNA/DNA ligation product produced over time.

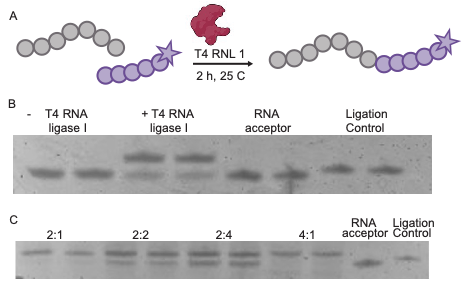

**Supplementary Figure 4.** Ligation of modified signal enhancer oligonucleotide to RNA acceptor (A) Ligation schematic (B) SYBR gold stained polyacrylamide gel electrophoresis of RNA acceptor with alkyl modified DNA oligonucleotide (C) Ligation of RNA acceptor and unmodified DNA donor at various ratios of donor to acceptor.

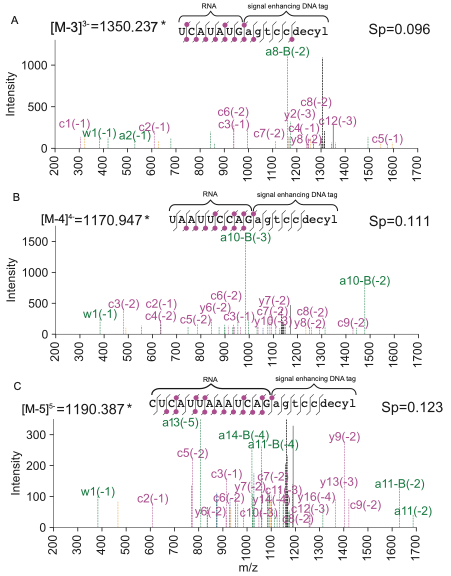
**Supplementary Figure 5.** Tandem mass spectra for RNA standard ligation products containing DNA signal enhancers at the 3′ terminus. RNA standards used had sequences UCAUAUG (A), UAAUUCCAG (B), and CUCAUUAAAUCAG (C). Only c- and y-ions are shown on the sequence ladder for clarity.

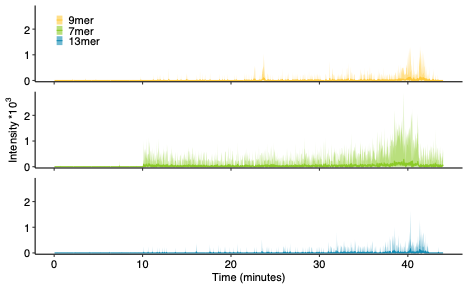

**Supplementary Figure 6.** Extracted ion chromatograms for RNA acceptors after ligation. The mass to charge ratios for the two most abundant charge states and corresponding sodium and potassium adducts are shown for the (A) 9mer (B) 7mer (C) 13mer acceptors.

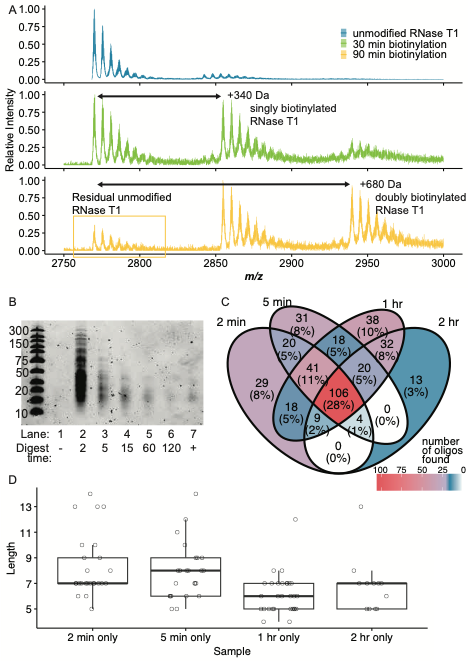

**Supplementary Figure 7:** (A) Mass spectra of RNase T1 before and after derivatization with NHS-LC-Biotin. Residual unmodified RNase T1 remains after 90 min reaction. (B) Polyacrylamide gel electrophoresis of 23S rRNA from *E. coli* digested with biotinylated T1 for various lengths of time. (C) Venn diagram of unique oligonucleotides identified in each sample using Pytheas. (D) Length distribution of unique oligonucleotides found exclusively in each sample.

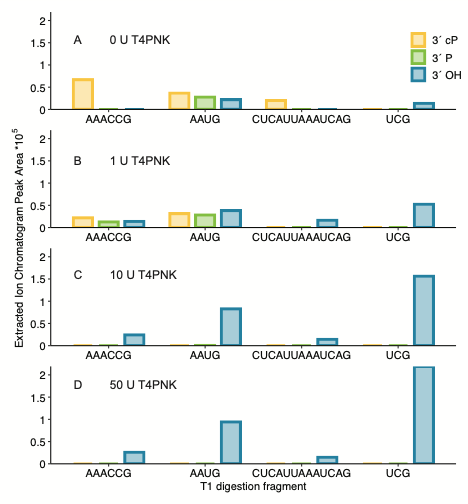

**Supplementary Figure 8.** Effect of T4 Polynucleotide Kinase on 3′ chemistry as measured by LC-MS extracted ion chromatogram peak area (A) 0 U T4 PNK, (B) 1 U T4 PNK, (C) 10U T4 PNK, (D) 50U T4 PNK.

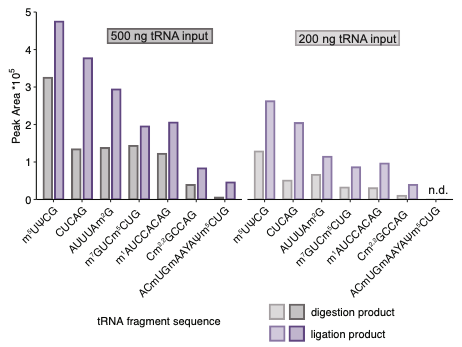

**Supplementary Figure 9:** Extracted ion chromatogram peak area comparisons between differing amounts of tRNA-Phe input. The fragment corresponding to the anticodon stem loop was not detected when 200 ng of input was used. Ψ signifies pseudouridine, Y signifies wybutosine.

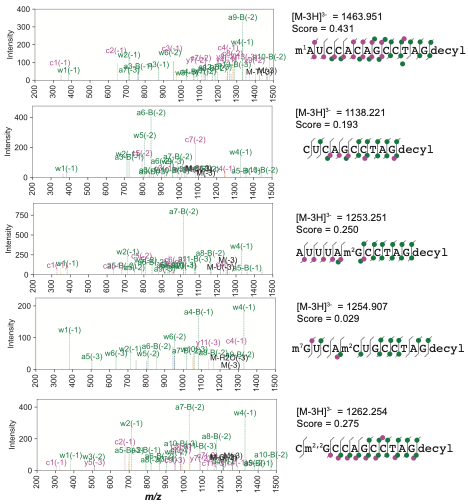

**Supplementary Figure 10:** Annotated tandem mass spectra from Pytheas for ligation products derived from tRNA-Phe.
